# *Pituitary carcinoma*: The University of Texas MD Anderson Cancer Center experience

**DOI:** 10.1101/411132

**Authors:** Fernando Santos-Pinheiro, Marta Penas-Prado, Carlos Kamiya-Matsuoka, Steven G Waguespack, Anita Mahajan, Paul D Brown, Komal B Shah, Gregory N Fuller, Ian E McCutcheon

**Affiliations:** Department of Neurology. Medical College of Wisconsin. 9200 W Wisconsin Ave. Milwaukee, WI 53226.; Department of Neuro-oncology. MD Anderson Cancer Center. Houston, TX.; Department of Endocrinology. MD Anderson Cancer Center.; Department of Radiation Oncology. Mayo Clinic, Rochester, MN.; Department of Radiology. MD Anderson Cancer Center. Houston, TX.; Department of Pathology. MD Anderson Cancer Center. Houston, TX.; Department of Neurosurgery. MD Anderson Cancer Center. Houston, TX.

**Keywords:** pituitary carcinoma, neuroendocrine tumors, temozolomide, chemoradiation.

## Abstract

*Background:* Pituitary carcinoma (PC) is an aggressive neuroendocrine tumor diagnosed when a pituitary adenoma (PA) becomes metastatic. PCs are typically resistant to therapy and frequently recur. Recently, treatment with temozolomide (TMZ) has shown promising results, although the lack of prospective trials limits accurate assessment. *Methods:* We describe a single-center experience in managing PC over a 22-year period and review previously published PC series. *Results:* 17 patients were identified. Median age at PC diagnosis was 44 years (range 16-82), and the median PA-to-PC conversion time was 5 years (range 1-29). Median follow-up was 28 months (range 8-158) with 7 deaths. Most PC were hormone-positive based on immunohistochemistry (n=12): ACTH (n=5), PRL (n=4), LH/FSH (n=2), GH (n=1). All patients underwent at least one resection and one course of radiation after PC diagnosis. Immunohistochemistry showed high Ki-67 labeling index (>3%) in 10/15 cases. Eight patients (47%) had metastases only to the CNS, and 6 (35%) had combined CNS and systemic metastases. The most commonly used chemotherapy was TMZ, and TMZ-based therapy was associated with the longest period of disease control in 12 (71%) cases, as well as the longest period from PC diagnosis to first progression in 8 (47%) cases. The 2, 3 and 5-year survival rate of the entire cohort was 71%, 59% and 35%, respectively. All patients surviving >5 years were treated with TMZ-based therapy. *Conclusions:* PC treatment requires a multidisciplinary approach and multimodality therapy including surgery, radiation and chemotherapy. TMZ-based therapy was associated with higher survival rates and longer disease control.

**Precis:** We describe 17 PC patients who were diagnosed and treated at MDACC over a 22-year period. We have found that TMZ-based therapy correlated with longer disease control and higher survival rate.

## Introduction

Pituitary carcinoma (PC) is a rare and aggressive neuroendocrine tumor (NET) accounting for approximately 0.1% of all pituitary neoplasms.^1^ The diagnosis is established after a pituitary adenoma (PA) becomes metastatic.^2-4^ Although metastases along the neuro-axis are more frequently observed, spread outside the CNS is also seen. The typical PA to PC latency period is several years, and little is known of the drivers for dissemination. Although p53 expression and Ki-67 labeling index correlate well with the degree of peritumoral invasion and aggressive behavior in PA, no specific histological or molecular markers are required to diagnose PC.^3, 5^ The management is challenging and a combination of surgical resection and/or radiation therapy is typically recommended, with chemotherapy often used when surgery and radiation are not possible or were previously unsuccessful. Nevertheless, response to standard treatment is usually transient and PC recurrence is common, with a reported median overall survival (OS) of 1 year.^1, 6^ Improved outcome has been recently reported with the use of chemotherapy in recurrent pituitary neuroendocrine tumors,^1, 7^ especially temozolomide (TMZ)^8-18^ as a single agent, in combination with capecitabine^19^ or concurrently with radiotherapy^20, 21^. Recent practice guidelines for the management of aggressive pituitary tumors and PC have recommended temozolomide as first line chemotherapy.^22^ Here we summarize a 22-year single-center experience treating newly diagnosed and recurrent PC, with an emphasis on the use of TMZ-based therapy.

## Methods

We conducted a retrospective review of all adult PC patients included in the University of Texas MD Anderson Cancer Center (MDACC) institutional databases from October 1, 1994 through January 31, 2017 under a protocol with waiver of consent approved by the Institutional Review Board. All patients had undergone a biopsy or surgical resection of a pituitary mass and/or metastatic disease. Patients’ demographic and clinical characteristics, treatments and outcome were reviewed. The diagnosis of PC was based on radiographic studies demonstrating metastatic dissemination and confirmed in equivocal cases by diagnostic biopsy or resection of a CNS or systemic lesion. Tumors were classified based on immunohistochemistry (IHC) findings and blood hormone levels (when available) into somatotroph (GH-PC), lactotroph (PRL-PC), corticotroph (ACTH-PC), thyrotroph (TSH-PC), gonadotroph (LH/FSH-PC) and null cell (NC-PC) subtypes.^23^ Since each PC patient received several lines of treatment (surgery, XRT, chemotherapy), we calculated the period of disease control achieved by each treatment modality. Disease control was defined as the presence of stable or decreased tumor burden in the primary sellar/locally invasive tumor and at all metastatic locations. We then determined which treatment modality was associated with the longest period of disease control. We calculated overall survival from the initial diagnosis of PC to death. PC progression was defined based on radiographic evidence of new or interval growth of metastatic lesions or of the primary tumor itself. Death was confirmed by review of medical records and/or death certificate.

## Results

### Demographics

A total of 17 patients with PC were seen over the study period. There was no gender predilection (Male:Female = 9:8, Table 1). The median age at diagnosis of PC was 44 years (range 16-82 years). Thirteen patients with PC were diagnosed and treated at MDACC, while 4 had the PC diagnosis confirmed in our institution but were treated elsewhere. Of note, 3 of the study subjects have already been presented in previous publications.^21, 24^

**Table 1:** PC, pituitary carcinoma; NC, null cell; ACTH/PRL/LH/FSH/GH, hormone-secreting PC; PA, pituitary adenoma; SRS, stereotactic radiosurgery; RT+TMZ, concurrent chemoradiation; RT, photon radiotherapy; PRT, proton radiotherapy; TMZ, temozolomide; cispl, cisplatin; CAPTEM, capecitabine+temozolomide; etop, etoposide; carbo, carboplatin; 5FU, 5-fluorouracil; PD-1, PD-1 inhibitor; pacl, paclitaxel; vinc, vincristine; CyVADIC, cyclophosphamide+vincristine+doxorubicin+ dacarbazine; bev, bevacizumab; CSRT, craniospinal radiotherapy; LMD, leptomeningeal disease; *, therapy prior to longest PFS; †, previously published cases from MDACC in the literature.

| PC subtype | Age (y), Gender | PA to PC latency (y) | Therapy upon PC diagnosis | PC recurrences | Therapy upon PC recurrence | CNS metastasis | Extra-CNS metastasis | Current status | Longest PFS (y) | OS (y) |
| --- | --- | --- | --- | --- | --- | --- | --- | --- | --- | --- |
| NC | 46, M | 8 | TMZ* | 1 | - | dura, LMD | - | deceased | 2.5 | 3 |
| NC | 44, F | 1 | RT+TMZ→cispl* | 0 | - |  | - | surveillance | 2.5+ | 3+ |
| NC | 69, M | 9 | TMZ* | 1 | CSRT | dura | bone | beva | 3.5 | 4+ |
| NC† | 69, F | 15 | RT+TMZ→CAPTEM* | 0 | - | - | lymph node, bone | surveillance | 3.5+ | 4+ |
| NC | 36, M | 11 | none* | 4 | TMZ | dura | lung, bone | TMZ | 13 | 13.5+ |
| ACTH | 45, M | 2 | RT→cispl+etop* | 0 | - | - | liver | deceased | <0.5 | <1 |
| ACTH | 54, F | 29 | RT→TMZ* | 0 | - | dura, LMD | - | surveillance | 2+ | 2+ |
| ACTH† | 41, M | 4 | RT | 10 | TMZ*, CAPTEM, CCNU+bev | dura | liver, orbits, lymph node | immunotherapy | 1.5 | 5.5+ |
| ACTH | 30, F | 11 | RT | 4 | SRS, TMZ* | dura, LMD | bone | TMZ re-challenge | 3+ | 6+ |
| ACTH† | 16, F | 5 | RT | 3 | CAPTEM, carbo-5FU* | dura | - | surveillance | 2 | 10.5+ |
| PRL | 81, F | 12 | RT | 2 | TMZ* | dura | lung | deceased | <1 | <1 |
| PRL | 47, M | 2 | none | 0 | - | dura | - | deceased | <0.5 | <1 |
| PRL | 25, M | 2 | RT→TMZ* | 0 | carbo+pacl, CAPTEM | dura | bone, liver | deceased | 2.5 | 10.5 |
| PRL† | 57, F | 6 | cispl+etop | 4 | TMZ*, CAPTEM, PD-1 | - | bone, liver | immunotherapy | 1.5 | 2.5+ |
| LH/FSH | 23, F | 2 | CyVADIC* | 1 | CyVADIC | dura | - | deceased | <1 | 1.5 |
| LH/FSH | 46, M | 4 | TMZ* | 4 | XRT+TMZ, RT | epidural spine | - | deceased | <1 | 5 |
| GH | 23, M | 2 | TMZ* | 0 | - | dura | - | TMZ | <0.5+ | <1+ |

### PA to PC transformation

The median PA to PC time to transformation (TTT) was 5 years (range 1-29 years, Table 1). One patient was diagnosed simultaneously with PA and metastatic disease (*de novo* PC). Among the IHC hormone-positive subtypes, ACTH-PC had the longest median TTT (5 years, range 2-29), followed by PRL-PC (4 years, range 2-12 years), LH/FSH-PC (3 years, range 2-4 years) and GH-PC (2 years). The median TTT of NC-PC was 9 years (range 1-15 years).

All patients were symptomatic at the time of diagnosis of PC and clinical presentation most commonly involved neurological symptoms such as headaches (n = 9 [53%]) and visual impairment (n = 5 [30%]). Other symptoms such as facial pain, focal weakness, increased thirst, and weight gain were also reported.

### PC subtype and molecular characterization

Pituitary tumors were classified based on IHC findings into somatotroph (GH-PC), lactotroph (PRL-PC), corticotroph (ACTH-PC), gonadotroph (LH/FSH-PC) and null cell (NC-PC). IHC was not available in 2 cases which were therefore classified based on blood hormone levels (1 PRL-PC and 1 GH-PC); there were no cases of TSH-PC, plurihormonal, or double tumors. The majority of PC (n=12) were hormone-positive: ACTH-PC (n=5), PRL-PC (n=4), LH/FSH-PC (n=2) and GH-PC (n=1). Five patients were NC-PC (Table 1). The one GH-PC patient had clinical signs of acromegaly. Three out of the 5 ACTH-PC patients developed Cushing syndrome.

Either before or after PC diagnosis was suspected based on imaging findings of metastatic lesions, all patients underwent at least one surgical resection of the pituitary tumor (mean of 2.3, range 1-6). IHC studies (15 sellar tumor samples available) showed a median Ki-67 index of 11% (range 1-40%); high Ki-67 labeling index (>3%) was present in 10/15 (67%) sellar tumor samples. Staining for mutant P53 was positive in 2 out of 9 tumor samples tested. Assessment for MGMT expression was not available in this cohort as it was not routinely done as part of clinical care.

Molecular studies via PCR-based next generation sequencing of tumor samples (primary and/or metastatic site) was done in 5 patients, and a hypermutator phenotype was present in 3 of them (ACTH-PC=2; NC-PC=1): in one ACTH-PC patient, an initial 46-gene panel was negative, but when a 400-gene panel was used, 20 potentially actionable gene mutations were identified (*ATM, GNAS, APC, CCNE1, CSF1R, EML4, ERBB3, ERBB4, FBXW7, FGFR4, IL7R, JAK1, MLL, MLL2, MLL3, NF2, NTRK3, PDGFRB, PTPN11, TOP1*) along with 53 other gene mutations identified in a tumor sample from tumor extension into the orbit. The other ACTH-PC patient had 1 potentially actionable mutation (*CREBBP* splice site *2086_2113+49del77*) along with another 20 gene variations of unknown significance identified in progressive sellar tumor. In the NC-PC patient, a metastatic tumor sample tested positive for MSH6; the MGMT methylation profile was negative. Among the two patients without hypermutator phenotype, a PRL-PC had no mutations identified in 147 genes tested in a metastatic tumor sample, while a NC-PC patient had a primary tumor sample positive for *FGFR1* amplification.

### Tumor location

Prior to PC diagnosis, 15 out of 17 pituitary tumors were locally invasive (one case had no information available about tumor extension prior to PC diagnosis; one case was diagnosed simultaneously with a PA and a distant CNS lesion). The most common site of local invasion was the cavernous sinus (n=8, 47%) and sphenoid sinus (n=8, 47%). Involvement of optic structures (optic nerves, orbit, chiasm) was observed in 35% of the patients (n=6), resulting in visual loss in 3 cases caused by compression of nerve rather than frank invasion. Most patients (n=13) developed central nervous system (CNS) metastases (dural based, n=13; leptomeningeal, n=3) (Figure 1A, 1B). In eight patients (47% of the entire cohort) metastasis was found only in the CNS, although the systemic work-up was not standardized in this cohort. Six (35%) patients had concomitant CNS and extra-CNS metastases (bone, n=3; lymph nodes, n=1; other organ (liver, lung), n=2) (Figure 1B), with two of them having more than one location of extra-CNS metastasis. Three (17%) patients had extra-CNS metastases without any metastases within the CNS (Table 1).

**Figure 1.**
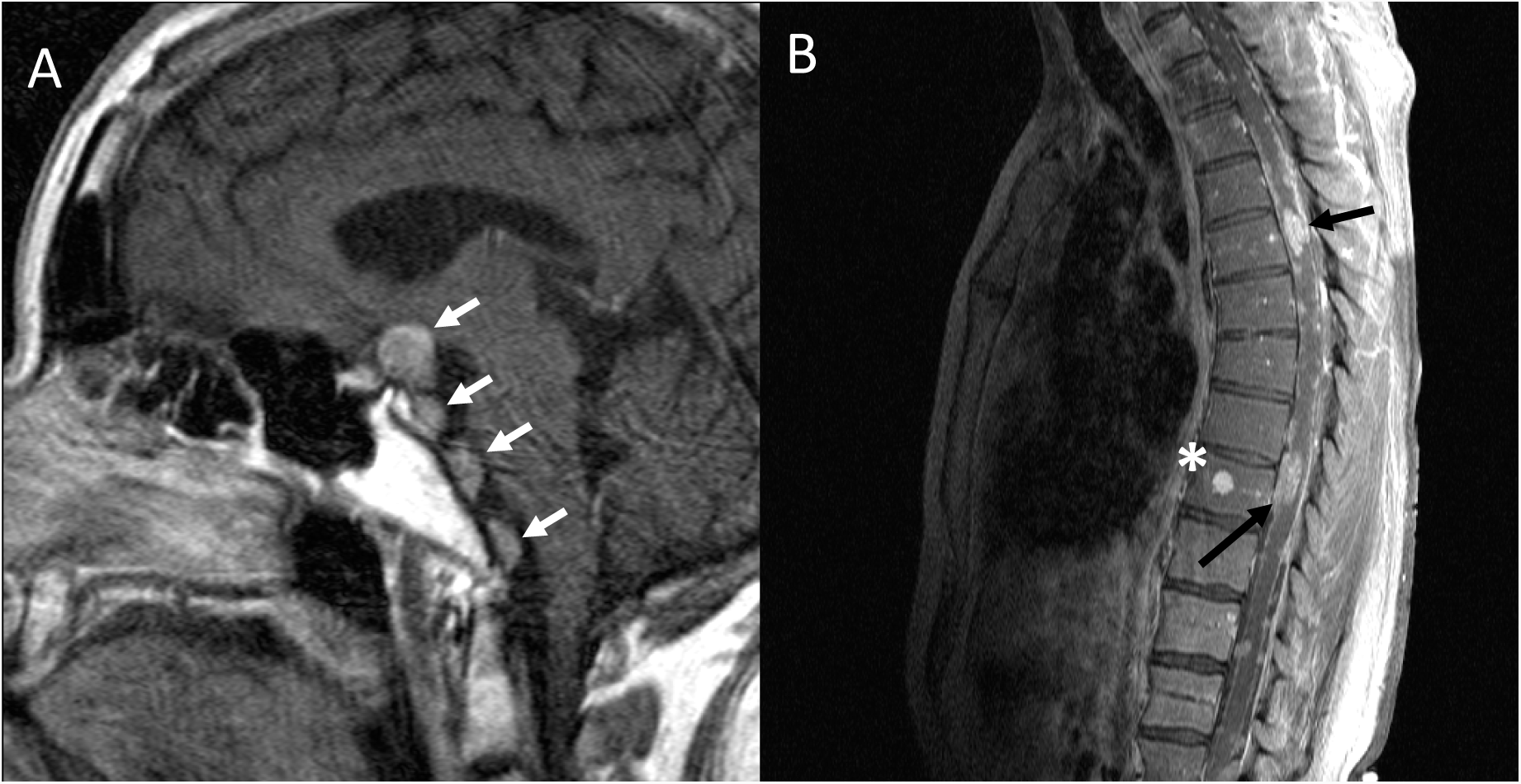
A) 17 year old girl with atypical pituitary adenoma, previously treated with surgery and radiation, presented with Cushing syndrome. Sagittal T1 post-contrast image shows multiple dural-based metastases (white arrows). B) 48 year old man with surgically proven dural-based metastases from pituitary carcinoma was treated with multiple surgeries, radiation and chemotherapy. He presented with foot drop. Sagittal T1 post-contrast fat-saturated image shows presumed bone metastasis (*) and innumerable intrathecal metastases (black arrows).

### Treatment modalities and outcome

The majority of patients received multimodality therapy.

### 4.1 Surgery

The average number of surgical pituitary resections from the initial PA diagnosis was 1.8 per patient (range 1-3). Seven patients (41%), required a craniotomy for further resection after an initial transsphenoidal sub-total resection of PA. The most common surgical techniques for resection of pituitary tumor were endonasal transsphenoidal approach (n=14) (including 1 endoscopic approach), followed by) pterional craniotomy (n=10) and sublabial transsphenoidal approach (n=2). Other surgical approaches consisted of transfacial/ lateral rhinotomy and fronto-orbito-zygomatic craniotomy (1 case each). Complications related to surgery occurred in 2 patients and consisted of cerebrospinal fluid (CSF) leak following the fronto-orbito-zygomatic approach requiring redo craniotomy and packing of the frontal sinus and sella, and repositioning of a ventriculoperitoneal shunt (which was placed following prior surgical resection of a PA that was causing hydrocephalus) following redo transsphenoidal approach for subsequent pituitary tumor resection. Of note, 3 ACTH-PC patients underwent bilateral adrenalectomy prior to PC diagnosis to control refractory Cushing’s disease.

### 4.2 Radiation Therapy

Following PC diagnosis, 7(41%) patients were treated with radiotherapy to the sella (n=5) or radiotherapy concurrent with TMZ (n=2). Intensity-modulated radiation therapy (IMRT, total dose: 45-54 Gy, divided in 25-30 fractions) was the most common radiotherapy modality in this setting. With regards to metastatic disease, a total of 13 foci were treated with radiotherapy (CNS metastases: n=7; extra-CNS metastases: n=6). IMRT was the most frequent technique (n=10), followed by stereotactic radiosurgery (SRS, n=4), craniospinal radiation (CSRT, n=2, both in the setting of leptomeningeal carcinomatosis) and intensity-modulated proton therapy (IMPT, n=1).

#### 4.3 Chemotherapy

TMZ was the most frequently prescribed chemotherapy in this series (Table 1), and it was used in all PC subtypes: NC= 5/5; ACTH= 4/5; PRL= 3/4; LH/FSH= 1/2; GH= 1/1. TMZ was used for newly diagnosed PC as monotherapy (n=4), concurrently with radiotherapy (n=2) and immediately following radiotherapy (n=2). The most common TMZ scheduling and dosage was 150mg/m2 for 5 days on a 28-day cycle, for 12 months (n=6). TMZ was also used for recurrent PC in 7 patients; it was given as monotherapy (n=5), combined with capecitabine (n=2; CAPTEM: capecitabine 1200 mg/m2/day in divided doses twice daily, days 1 to 14, and TMZ 150-200 mg/m2/day in divided doses twice daily, days 10 to 14 of a 28-day cycle, for a total of 6 cycles)^11^, or concurrently with radiotherapy (n=1; TMZ 75 mg/m2 daily for 42 days combined with radiation 5 days a week for 6 weeks). Four patients (one LH/FSH-PC, one ACTH-PC and two PRL-PC) were re-challenged with a TMZ-based therapy upon PC recurrence. One of them (LH/FSH-PC) was treated with TMZ upon PC diagnosis and had recurrence less than a year later; this patient then received concurrent TMZ and radiotherapy and died less than one year later (OS 5 years). The other 3 patients were treated with TMZ-based therapy upon first and subsequent PC recurrences; all of them were initially treated with TMZ alone (which conferred a median PFS of 1.5 years, range 1-2.5 years), and were then treated with CAPTEM, which provided progression-free survival of less than 1 year.

TMZ therapy was overall well tolerated with anticipated side effects, and in none of the patients was the therapy terminated or delayed due to hematological or non-hematological adverse events. The most common adverse events attributed to TMZ were fatigue and nausea (35%), with no reported grade III-V toxicity, according to CTCAE v4.0 (2009).

Other chemotherapy regimens used for newly diagnosed PC were cisplatin (after concurrent radiation and TMZ, n=1), cisplatin plus etoposide (n=1), and cyplophosphamide in combination with vincristine, doxorubicin and dacarbazine (CyVADIC, n=1). In the recurrent setting, treatments included bevacizumab as a single agent (n=3), bevacizumab plus irinotecan (n=1), bevacizumab plus pan-FGFR kinase inhibitor (n=1, as part of a phase 1 clinical trial), bevacizumab plus lomustine (n=1), carboplatin plus vincristine (n=1) and carboplatin plus 5-FU (n=1)^24^.

In this series, 3 patients were enrolled in 3 different phase 1 clinical trials testing: 1) tipifarnib plus sorafenib, a farnesyltransferase inhibitor and a multi-tyrosine kinase inhibitor (VEGFR, PDGFR, *Raf*), respectively; 2) pembrolizumab, a PD-1 inhibitor, and 3) BGJ398, a pan-FGFR inhibitor. The outcome of these patients treated with experimental drugs is not discussed here.

#### 4.4 Outcome

Progression after any given therapy, i.e., after first treatment for newly diagnosed PC and after any treatment for subsequent progression, if applicable, occurred in 10 (59%) patients, at a median time of 18 months (range 0.33 - 13 years) and at a median occurrence of 1 (range 1-10 occurrences) per patient. Of the 7 patients without imaging evidence of recurrence during follow up, 4 were diagnosed with PC less than 3.5 years prior to the end of the study and were still alive, and 3 patients were deceased in less than one year from PC diagnosis. The cause of death was unknown, but death due to progression of PC could not be excluded. Median time from PC diagnosis to first PC recurrence was 9 months (range 3->42 months). Eight patients (47%) were treated with upfront TMZ or TMZ-based therapy (Table 1) and achieved a median time from PC diagnosis to first PC recurrence of 30 months (range 5->42 months). Another 8 patients (47%) were treated with upfront chemo- or radiotherapy without TMZ and achieved a median time from PC diagnosis to first PC recurrence of 10 months (range 0.33-13 years). One patient (PRL-PC) received no treatment due to poor performance status at PC diagnosis (OS 8 months). One patient (NC-PC, initial metastatic site: cerebellum) achieved 13 years of stable disease without upfront therapy and is now on TMZ following the 4^th^ recurrence of disease (further metastatic sites: pons, lung and bone). Overall, TMZ-based therapy was associated with the longest period of disease control in 12 (71%) patients when used at diagnosis or recurrence (median 21 months, range 4->42 months, Table 1). The median follow-up after PC diagnosis was 2 years (range 1-13 years). The 2-, 3- and 5-yearsurvival rates for the entire cohort were 71% (n=12), 59% (n=10), and 35% (n=6), respectively. Of the patients who were alive at least 5 years from diagnosis, 2 patients (one NC-PC, one ACTH-PC, both still alive) survived more than 10 years and all 5 had received treatment with TMZ-based therapy (Table 1). One refractory case is currently under treatment with the PD-1 inhibitor pembrolizumab as part of a phase 1 clinical trial. Eighty percent of NC-PC were alive at last follow-up, in contrast with 58% of IHC hormone-positive PC. There were 7 reported deaths by the end of the study period. Cause of death was infectious etiology in 1 patient (pneumonia, while off temozolomide for one month) and unknown in 6 patients.

## Discussion

Pituitary carcinoma (PC) is a rare and aggressive form of pituitary tumor for which limited therapeutic guidelines exist. As no curative treatment has been established for PC to date, this case series aimed to present our experience with the use of multiple modalities of treatment over a prolonged period of follow-up (median, 2 years; range, 1-13 years). Our experience suggests that a multimodality approach involving a combination of surgical resection, radiotherapy and chemotherapy (particularly TMZ-based) correlates with better outcome (2-year OS rate 71%, 3-year OS rate 59%, 5-year OS rate 35%), compared with previous reports in the literature (Table 2).^1, 6, 7, 21^

**Table 2:** Previously published pituitary carcinoma case series by single and multicenter groups. *: median PFS and OS since PC diagnosis. Abbreviations: PC, pituitary carcinoma; TMZ, temozolomide; PFS, progression-free survival; O, overall survival; ACTH, adrenocorticotrophic hormone; PRL, prolactin hormone; NC, null-cell; GH, growth hormone; FSH/LH, follicle-stimulating hormone/luteinizing hormone; NOS, not otherwise specified; RT, radiotherapy;

| Study | Study period | Single center | PC cases | PC type | TMZ Tx | other Tx modalities | PFS*(mo) | OS*(mo) |
| --- | --- | --- | --- | --- | --- | --- | --- | --- |
| *Pernicone, 1997*<sup>7</sup> | 1955-1994 | Yes | 15 | ACTH(7), PRL(7), NC(1) | 0 | surgery, RT, cytotoxic agents | Not reported | 1-y OS 66%, mOS 22 |
| *Kaltsas, 1998*<sup>41</sup> | 1970-1997 | Yes | 4 | ACTH(1), PRL(3) | 0 | cytotoxic agents, RT | 11.5 | 11.5 |
| *Roncaroli, 2003*<sup>42</sup> | 1983-2013 | Yes | 2 | PRL(2) |  | cytotoxic agents | 9 | 10 |
| *Raverot, 2010*<sup>43</sup> | Not reported | No | 5 | ACTH(2), PRL(3) | 5 | surgery, RT | Not reported | Not reported |
| *Hirohata, 2013*<sup>26</sup> | 2006-2011 | No | 12 | ACTH(3), PRL(3), NC(4), NOS(2) | 10 | Not reported | Not reported | Not reported |
| *Hansen, 2014*<sup>44</sup> | 1973-2008 | No | 7 | Not reported | 0 | surgery | Not reported | 1-y OS 57.1%, 2-y OS 28.6%, 5-y OS 28.5% |
| *Bengtsson, 2015*<sup>18</sup> | Not reported | No | 8 | ACTH(3), PRL(2), GH(3) | 8 | surgery, RT, cytotoxic agents | Not reported | Not reported |
| *Wang, 2015*<sup>45</sup> | 2006-2015 | Yes | 2 | ACTH(1), GH(1) | 0 | surgery, RT | 156 (ACTH)<br>22 (GH) | 178 (ACTH)<br>34 (GH) |
| *Yoo, 2018*<sup>46</sup> | 2018 |  | 2 | ACTH(2) | 2 | surgery, RT | 6 | 12+ |
| *Santos-Pinheiro, 2018* | 1994-2017 | Yes | 17 | ACTH(5), PRL(4), GH(1), FSH/LH(2), NC(5) | 15 | surgery, RT | 18 | mOS 36; 2-y OS 65%, 3-y OS 53%, 5-y OS 29% |

According to the World Health Organization (WHO) classification of pituitary tumors (2017)^4^, the presence of metastatic disease in the CNS or systemically suffices to designate a pituitary tumor as PC. However, this definition carries important limitations. This approach disregards intrinsic histopathologic or molecular features of the pituitary tumor and relies exclusively on the detection of metastasis. However, despite metastatic activity, the metastatic disease itself may remain devoid of typical malignant features (i.e. p53 expression, tumor invasion, high proliferation rate).

The strategic location of the pituitary in relation to other intracranial structures (internal carotid artery, dura, cavernous sinus) may facilitate the hematogenous, CSF and lymphatic spread of cancerous cells into the neuraxis and extra-CNS locations. Yet, metastatic disease occurs in only 0.1% of all PA cases and little is known of the biologic drivers for malignant behavior in these tumors. The majority of patients in our series had CNS metastasis with involvement of the meninges (n=11), which raises the question of the mechanism of spread of PC, either by CSF or hematogenously, given its close proximity to vascular structures as well as the meninges lining the sella. Extra-CNS metastases involving vertebral bodies, long bones, liver, lung, and lymph nodes were seen in over half the patients in our series.

The spectrum between PA and PC likely implies biologic differences beyond the presence of metastases, although these are unknown at present. The 2017 WHO classification aimed to improve tumor classification by removing the term “atypical PA” as a pituitary tumor subtype (PA with increased proliferation rate [Ki-67 labeling index > 3% and increased mitotic figures], p53 overexpression, local tumor invasion, and atypical morphologic features). However, the new WHO classification still recommends the use of the term “aggressive PA” (invasion pattern, rapid recurrence or resistant to therapy) to classify tumors deviating from a ‘benign’ behavior. Although the terms ‘aggressive’ and ‘invasive’ have been used interchangeably in the literature, such mixing blurs the distinctions among rapid tumor growth, invasion of adjacent structures, metastasis to distant sites, and resistance to treatment. Such distinctions are necessary for a biologically relevant definition of an “aggressive PA.” Moreover, despite evidence of correlation between histologic proliferation markers and clinical aggressiveness in PA, no specific cutoff was recommended for Ki-67 and mitotic index by the 2017 WHO classification, which does not provide clear guidelines for classification or treatment of such tumors. In clinical practice, however, the presence of these histologic markers is commonly incorporated in decisions about frequency of surveillance and in discussions with patients about the potential risk for recurrence.

In our series, the mean latency period from diagnosis of PA to PC transformation was 5 years (range, 1-29 years), which is in accordance with the literature.^1, 6^ Among the IHC hormone-positive PC subtypes, GH- and LH/FSH-PC had the shortest latency period, while PRL-PC, ACTH-PC, and NC-PC had the longest latency. However, given the small number of patients with each subtype in our series, we cannot determine with certainty if this holds true in the entire population of patients with PC.

The Ki-67 labeling index was assessed in the sellar tumor in the majority of our patients (n=15) and, before the PC diagnosis was established, it was high (more than 3%) in 67% of them. The present series and review of the literature support that the vast majority of PC arise from previously locally infiltrative tumors with high proliferation rate; however, a low Ki-67 does not eliminate the risk of malignant transformation. Other pathologic biomarkers such as mutant p53 expression, MGMT IHC or promoter methylation status, MSH6 IHC, and mutation profile were not systematically tested in our series, although they may be of clinical relevance for prediction of treatment response (*i.e*., to alkylating agents), prognostication, and/or stratification in clinical trials ^25-27^.

The rationale for the use of TMZ in PC is supported by several clinical and pharmacological studies in neuroendocrine and primary brain tumors, indicating its excellent blood-brain barrier penetration, in addition to its homogenous distribution in CNS and extra-CNS tissues.^12, 28^ Ramanathan *et al.* reported an objective response rate of 34% in a phase II study using dacarbazine (DTIC) intravenously as single-agent therapy for metastatic pancreatic neuroendocrine tumors (PNETs).^29^ TMZ, which is bioactivated into the same metabolite as DTIC (5-methyltriazenoimidazole-4-carboxamide, MTIC), has the advantage of undergoing spontaneously decarboxylation (and therefore bypassing hepatic activation)^30^. The relatively mild toxicity profile and ease of use (oral formulation) are also advantages of TMZ over other chemotherapies. TMZ works by inducing cell apoptosis or cell senescence in rapidly dividing cells and it is non-specific to any mitosis phase. Furthermore, TMZ demonstrated good response in primary brain tumors and previously untreated brain metastases when added to radiotherapy.^31^ In parallel, the relatively low MGMT expression seen in NET, including PA and PC, increases their sensitivity to alkylating agents such as TMZ.^32, 33^ The 2017 European guideline for pituitary tumors encourages testing of MGMT expression by IHC on aggressive pituitary tumors and pituitary carcinomas (even though only a minority of them exhibit homogenous MGMT expression), as it may help predict treatment success with temozolomide, the first-line chemotherapy recommended for such tumors.^22, 34^ Nevertheless, MGMT may not be the sole driver for susceptibility to TMZ, and other enzymes, such as MSH6, may also contribute to PC sensitivity to alkylating agents.^25^ For this reason, caution is warranted when using MGMT expression as the only criterion to decide on the use of temozolomide in PC and locally aggressive PA, as both lack of response in patients with low expression and favorable response in patients with high expression have been described. ^35^ Therefore, a trial of therapy with TMZ may be warranted regardless of MGMT status, particularly if other treatment options have been exhausted.

Further reports described the effective use of TMZ combined with capecitabine in recurrent and metastatic PNETs^11, 12, 21, 36-38^. The addition of capecitabine to TMZ (CAPTEM) is based on the theory that sequential pretreatment with capecitabine may potentiate the cytotoxicity of TMZ by synergistically depleting thymidine, leading to apoptosis.^11, 28^ A recent randomized Phase II study has shown improved PFS and OS of capecitabine in combination with TMZ compared to TMZ alone in advanced pancreatic NETs;^39^ however, this has not been tested in PCs and whether CAPTEM is superior to TMZ alone in this patient population remains unknown. Potential benefits of using combined therapy must be carefully weighed against the potential for added toxicities. In our series, the use of CAPTEM therapy in patients with previous progression after TMZ monotherapy was limited to 3 cases, and definitive conclusions about the benefit of CAPTEM in the setting of TMZ failure cannot be drawn.

In our patient cohort, we observed a longer median time between PC diagnosis and first PC recurrence in patients treated with upfront TMZ-based therapy (30 months) compared with patients treated with other lines of upfront therapy that was not TMZ-based (10 months), although the low number of patients and the potential bias inherent in retrospective series limits generalization of the validity of these results. Our results are also superior to those previously reported in the literature (Table 2); this includes results from a contemporary study in which TMZ was commonly prescribed, and which reported that 42.5% of PC patients were deceased with median follow-up of 12 months.^40^ Nevertheless, we are unable to confirm whether the favorable outcome seen with TMZ is due to “first therapy effect”, when the longest PFS is usually seen, whether it is related to higher TMZ efficacy or, more likely, whether it is a combination of both.

Radiotherapy with concurrent TMZ was completed in 3 patients, which was then followed by cisplatin (n=1), TMZ (n=1) and capecitabine plus TMZ (n=1). Two of these patients had disease control for more than 2 years, and all 3 patients survived for at least 3 years. This is in agreement with a recent European survey which showed that concurrent TMZ and RT was associated with increased response rate compared to TMZ alone.^40^ The longer survival rates in our cohort may be correlated to the multimodality approach and/or TMZ use, especially considering that all long-term (>5 years) survivors were treated with TMZ-based therapy.

This study has the limitations typical of an observational retrospective study, including the small cohort and single-center setting. Therefore, we are unable to draw any comparative statistical analysis among the different treatment modalities given to each PC patient or across PC patients. Furthermore, a tertiary institution serving as a national and international referral center invariably has a selection bias for patients with more severe conditions and poorer prognosis. Finally, the confounding bias related to several lines of therapy in a relatively short timeframe also limits the interpretation of a true cause-effect phenomenon between each line of therapy and the observed outcome. Despite all these limitations, our series provides valuable information on the demographics, tumor characteristics, treatment modalities and outcome of patients with this rare malignancy for which prospective data are lacking to guide estimation of prognosis and decisions on therapy.

## Conclusion

PC is a rare and aggressive neuroendocrine malignancy. Local recurrence is frequent and metastases occur in both CNS as well as extra-CNS locations. A combination of surgical resection, radiotherapy, and chemotherapy (particularly TMZ-based) may result in prolonged survival. In this case series, which is the largest single-institution experience published in the literature, early use of chemotherapy, specifically TMZ, combined with standard PC management (surgical resection and radiotherapy) was well tolerated and associated with improved survival rates compared to the previous literature.

